# MCbiclust: a novel algorithm to discover large-scale functionally related gene sets from massive transcriptomics data collections

**DOI:** 10.1101/075374

**Authors:** Robert B. Bentham, Kevin Bryson, Gyorgy Szabadkai

**Author notes:** To whom correspondence should be addressed.; Correspondence may also be addressed to KB and GS.

## Abstract

The potential to understand fundamental biological processes from gene expression data has grown parallel with the recent explosion of the size of data collections. However, to exploit this potential, novel analytical methods are required, capable of handling massive data matrices. We found current methods limited in the size of correlated gene sets they could discover within biologically heterogeneous data collections, hampering the identification of multi-gene controlled fundamental cellular processes such as energy metabolism, organelle biogenesis and stress responses. Here we describe a novel biclustering algorithm called Massively Correlated Biclustering (MCbiclust) that selects samples and genes from large datasets with maximal correlated gene expression, allowing regulation of complex pathway to be examined. The method has been evaluated using synthetic data and applied to large bacterial and cancer cell datasets. We show that the large biclusters discovered, so far elusive to identification by existing techniques, are biologically relevant and thus MCbiclust has great potential use in the analysis of transcriptomics data to identify large scale unknown effects hidden within the data. The identified massive biclusters can be used to develop improved transcriptomics based diagnosis tools for diseases caused by altered gene expression, or used for further network analysis to understand genotype-phenotype correlations.

## INTRODUCTION

Present gene expression datasets typically contain thousands of samples, each measuring tens of thousands of genes. Moreover, the size of the currently generated sample-gene matrices continues to increase dramatically with the advances of more economical high throughput technologies. These extensive datasets hold the promise for the discovery of novel regulatory networks underlying fundamental physiological and pathological cellular processes governed by multitudes of genes, such as cellular energy and redox metabolism, organelle biogenesis and integrated stress responses (1–5). Indeed, while quantitative models of networks involving genes on relatively small scale have been now well established (see e.g. (6–9) for examples related to metabolism), bioinformatic discovery approaches capable of handling large datasets are in critical need of development.

Currently, extracting information on biological processes from genomic, transcriptomic and proteomic datasets relies on a pipeline including (i) identification of frequent genomic mutations or differentially represented transcripts or proteins, followed by (ii) pathway and network analysis methods using gene-set, pathway or network databases (for a recent reviews see e.g. (10)). A number of effective approaches for both stages of the analysis have been developed, but they have considerable limitations.

First, differential expression algorithms (11, 12) are used to filter experimental data to find genes with significant alterations, producing lists that can be sorted into biologically relevant groups using gene set enrichment analyses. Recent developments, such as gProfiler (13) or GSEA (14) extended the value of this approach by considering a ranked or continuous scale of gene expression differences, as opposed to methods using unranked sets of genes chosen with fixed gene expression p-value thresholds (e.g. DAVID (15)). However, interactions and potential co-regulation of genes are not considered in these approaches, thus they can only be used to assign previously determined fixed gene sets enriched in the data. Most functionally annotated pathways represent normal physiology, thus the use of these methods excludes the possibility to discover novel functional groups relevant to stress and pathologically relevant pathways. One approach to currently overcome this limitation is to apply methods incorporating databases with rich information on gene or protein interactions, such as BioGRID (16), IntAct (17), STRING (18) or GeneMANIA (19), and identifying networks with altered gene expressions. Numerous examples using this approach exist, such as GeneMANIA (19), ReactomeFIViz (20), STRING (18), ResponseNet (21), NetBox (22), MEMo (23) and EnrichNet (24). Whilst these approaches were proven successful in identifying altered core pathways in several pathologies, since they are based on prior knowledge of network components and structures, they have limited potential to discover novel co-regulated large scale networks determining cellular phenotypes. In this paper we argue that large scale differences in gene expression, for instance between different physiological and pathological states, go undiscovered due to these limitations and that novel methods discovering large-scale gene correlations are needed.

Another difficulty is that the now common large datasets used for network discovery are typically not produced by experimental design based on a priori knowledge but are more often mass data collection projects containing vastly heterogeneous samples. In many cases it is unclear how to divide these samples into subclasses, due to the many unknown factors distinguishing subtypes with different gene expression patterns. Hierarchical clustering has notably been used to find related subgroups of samples, notably first by Eisen (25) but also by Perou (26) who used this technique to identify the intrinsic subtypes of breast cancer. These standard clustering techniques however are only useful at finding strong patterns within the data, since they cluster all the samples against all the genes or vice versa, risking to omit more global weaker patterns, due to high noise. Modes of gene regulation could be present in only a subset of samples, with genes being conditionally co-regulated only on specific cellular or environmental signals (27). With only a subset of samples having this regulation, standard clustering techniques would not detect this co-regulation in the noise of the data. Thus our second consideration for developing a method solving this problem and discriminating heterogeneous samples with co-regulated genes in large datasets was to use biclustering algorithms.

Biclustering techniques were first applied to gene expression by Cheng and Church (28), but the technique itself dates back to the 1970's in the work of Hartigan who referred to it as direct clustering (29). Biclustering algorithms select a subset of the rows and columns of a data matrix such that a particular measurement describing the quality of the bicluster is maximised. It is not known a priori how many significant biclusters there are within a data matrix, and the number and size of found biclusters depend on (i) the definition of bicluster, (e.g. correlation of gene expression across samples) (ii) the method of measuring its quality, and (iii) the method for searching for biclusters. There are a large number of existing biclustering algorithms involving different quality metrics as well as search heuristics for finding them (30), but we have found them of limited use for the scope of finding large co-regulated gene sets in a subset of samples within massive datasets. Mean square residue score for evaluating biclusters (28) is used in many biclustering techniques (MSB (31), FLOC (32), BiHEA (33), etc.). As a quality metric it does find biologically relevant biclusters but is limited to finding bicluster involving a simple shift in gene expression between samples but not patterns which involve more pronounced scaling of gene expression (34). Moreover, most of these methods are not computationally efficient on very large datasets, since finding biclustering has been shown to be an NP-hard problem (35), much more difficult than normal clustering. Accordingly, existing biclustering algorithms are adept at finding many small sized biclusters involving relatively few genes but not suitable for discovering large scale biclusters.

Here we describe the development of a conceptually novel biclustering algorithm, based on evaluating correlated gene expression across large sets of heterogeneous samples. The approach, in contrast to previous methods, is (i) computationally efficient to be applied to large data matrices containing whole genome transcriptomic data of more than a thousand samples, and (ii) capable to identify correlated, biologically relevant large gene sets and subsets of heterogeneous samples where the gene set is being differentially regulated. The method addresses key questions in genome biology. First, by quantifying correlations and expression levels of the discovered gene sets the method can be applied to classify samples. In addition, the gene sets can be used for discovery of large networks, controlled by master transcriptional regulators, which thus likely determine fundamental cellular phenotypes.

## MATERIAL AND METHODS

### The MCbiclust workflow

Massively correlated biclustering (MCbiclust) is a stochastic iterative search based method that uses Pearson’s correlation coefficient as a quality metric to find biclusters (Fig. 1).). The basic strategy is to start with around 1000 seed genes and a small number of seed samples, then through random replacement of samples find a bicluster that can be then expanded. MCbiclust is specifically designed to find biclusters composed of large numbers of genes and samples within data sets. The hypothetical ideal bicluster is one whose genes are highly correlated across all samples in the bicluster, and it is not important whether these correlations are positive or negative. The algorithm is stochastic and each run will end with a different massively correlated bicluster being discovered. So generally the method is run many times, typically up to a thousand, to discover the key large-scale biclustering structure within the given data collection. All the biclusters discovered are compared to determine how many different biclustering groupings exist.

**Figure 1.**
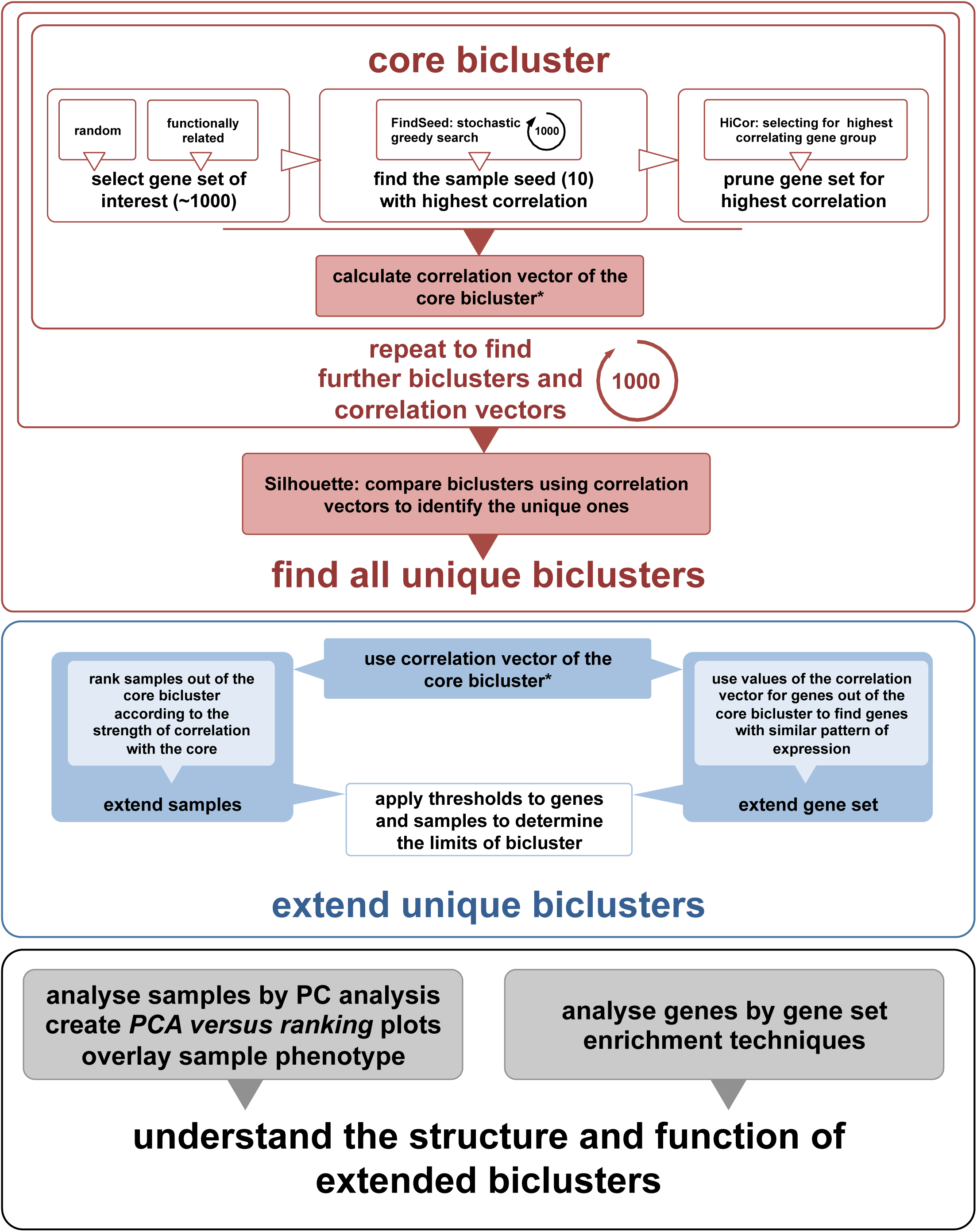
Schematic overview of the MCbiclust pipeline. The schematic shows (i) the methods used to find a core bicluster how this process is repeated and compared by Silhouette analysis to identify the unique biclusters (upper panel); (ii) how these biclusters are then extended (middle panel); and (iii) functionally and structurally analysed (lower panel). The overall description of the process is given in the Materials and Methods section, with full details of each step describes in Supplementary Methods. A key step in the bicluster analysis is the calculation of correlation vectors, which is further explained in Supplementary Fig. S1.

For each individual run, the algorithm starts with an initial seed of 1000 genes that are either chosen randomly for discovering general large-scale features in the data collection, or are chosen for functional relevance to direct the discovery of biclusters (for instance a mitochondrial related gene set). Each run starts with a random seed of 10 samples. A greedy search is then undertaken where individual samples are randomly replaced by other samples, with the aim of always increasing the overall correlation score of the bicluster. Once 10 samples have been determined that maximize the bicluster correlation score, the pipeline focuses on the genes involved to further maximize this score. Hierarchical clustering of the genes is carried out, dividing the genes into 8 groups with tightly correlated genes over the samples, only the genes from the group which has maximum bicluster score are kept with all the other genes being removed.

Now that the nucleus of a highly correlated bicluster has been formed, the bicluster is extended in terms of both samples and genes included. An “average gene expression vector” is determined from the bicluster, by dividing the genes into groups with hierarchical clustering and finding the average gene expression of this group across the 10 samples. The correlation of every gene measured to this average gene expression vector can be calculated forming a “correlation vector”. The genes can then be ordered by their values in the correlation vector (See Supplementary Fig. S1). Following gene extension, all the other samples within the data collection can be ranked according to how well they preserve the correlation of the bicluster. At each step the sample that preserves most the correlation is added, until all the samples have been ranked. MCbiclust therefore returns a ranked list of the samples and genes matching the pattern found in the bicluster. In order to determine which genes and samples are in the bicluster a method to threshold the bicluster is applied as described in Supplementary Methods.

The biclusters discovered are often complex and thus we have used two key approaches to interpret them in terms of either the samples or genes involved. Samples are analyzed by doing Principal Component Analysis (PCA) across gene values across the 10 most prominent samples. The first principal component (PC1) is then used to visualize each of the samples within the bicluster ranked according to correlation. Generally such plots split the samples into two forks with anti-correlated gene expression between two groups of genes identified (see Supplementary Fig. S3). The key approach employed to analyze the genes within a bicluster in order to help identify its biological nature is gene enrichment analysis. Although it can be seen later that bicluster interpretation often needs investigation driven by intuition based on considering both the samples and genes involved.

Detailed information about the algorithm can be found in Supplementary Methods and in the Vignette accompanying the Bioconductor package developed to perform custom MCbiclust analysis.

### Synthetic Data and Benchmarking

A synthetic dataset was created using an adapted version of the method used in (19) for the biclustering method FABIA, using the R package `FABIA’. This method implants a set number of multiplicative biclusters that match the FABIA model, into a dataset. This was done by creating 8 separate synthetic datasets, using the FABIA model. Each dataset contained only 1 bicluster, on average containing approximately 500 genes and 130 samples, and each dataset was mean centered according to the genes before being combined. Eight biclusters were chosen so that the final synthetic dataset contained 1000 genes and 1059 samples. Enforcing sample exclusiveness to a single bicluster was done primarily to make the comparison between the different bicluster algorithms feasible. If a sample belonged to two or more biclusters, due to each bicluster affecting a large number of the genes, there would be a significant number of genes belonging to both biclusters and this overlap of genes could potentially confound the classification of samples to their correct bicluster.

MCbiclust was compared with the FABIA (36), FABIAS (36), biMax (37), CC (38), Plaid (39), ISA (40), FLOC (41), QUBIC (42) and CPB (43) biclustering methods (See Supplementary Table 1). These methods were chosen due to their availability of access as R packages on bioconductor, or due to the quality metric similar to the one utilised in MCbiclust (CPB). CPB was run with a python script available at http://bmi.osu.edu/hpc/software/cpb/index.html.

### Workflow to compare biclusters obtained with different methods

Fig. 2A provides an overview of how the results of each biclustering method (shown as biclusters F1 to F8) were compared to the real biclusters present in the synthetic data (shown as A1 to Ax where x is the variable number of biclusters predicted). First, a similarity matrix is constructed where all possible predicted biclusters from the results are compared to all of the 8 known biclusters in the synthetic data. The Jaccard score is used since this is appropriate for comparing the similarity between two different sets (being equal to the number of elements in the intersection of the two sets divided by the number of elements in the union of the two sets). Identical sets will have a Jaccard score of 1.0 and completely different sets will have a Jaccard score of 0.0. Once all predicted biclusters are compared to all known biclusters in the matrix, the Hungarian or Munkres algorithm is used to efficiently determine the most optimal matching of predicted biclusters to known biclusters which maximises the sum of the scores (44). At this point each real bicluster (S1 to S8) would be matched to its most optimal predicted bicluster (A1 to A8) by the method. With this matching complete, traditional measurements of accuracy, false positive and true positive scores can be used both for the samples matched and the genes matched, and receiver operating characteristic (ROC) curves can be plotted.

**Figure 2.**
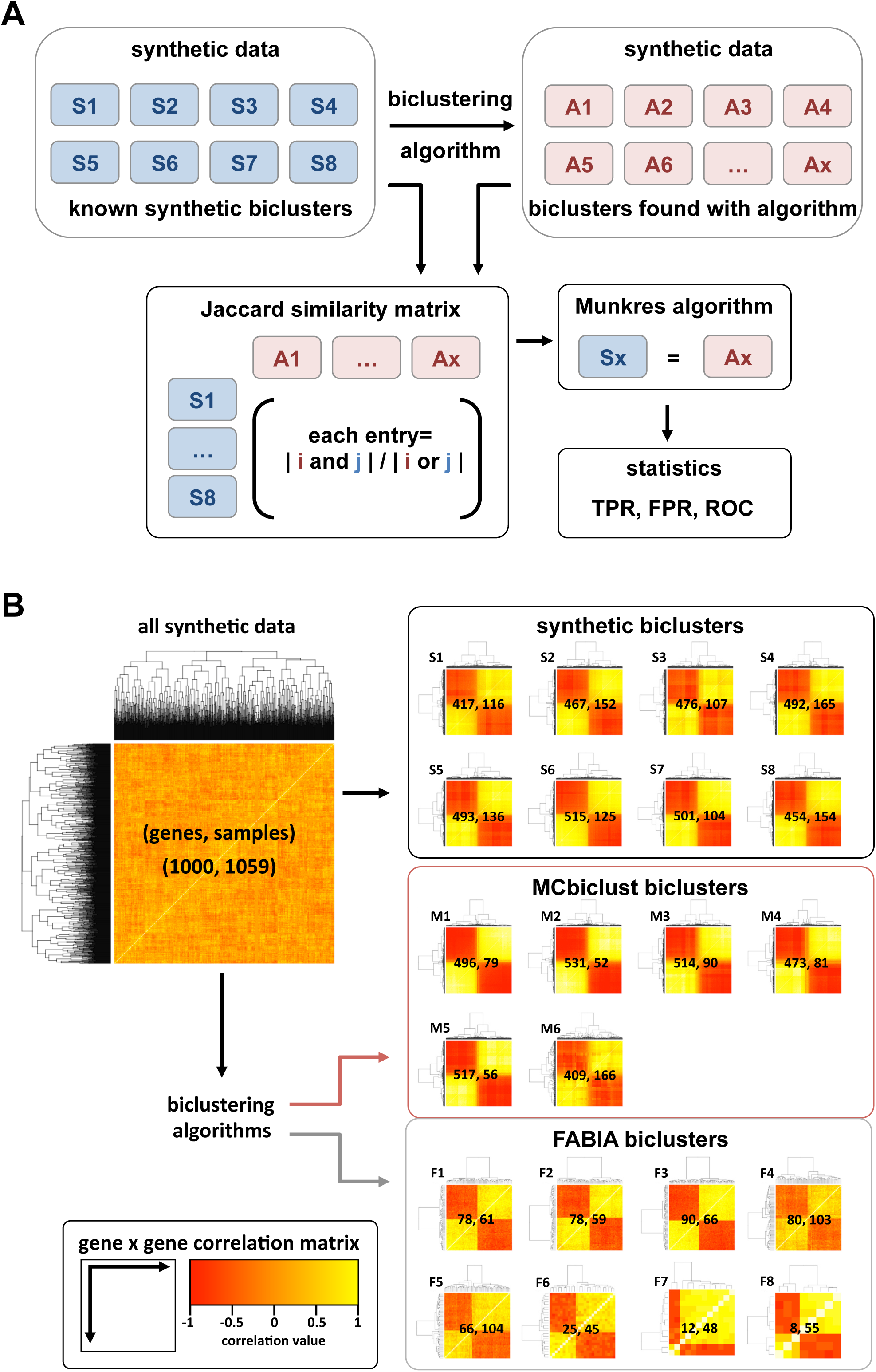
Benchmarking of MCbiclust against previous biclustering methods. **A.** Outline of the evaluation pipeline. Known biclusters in the synthetic datasets are compared with the biclusters found with different biclustering methods. Jaccard Index and the Munkres algorithm is used to solve the assignment problem of matching the known synthetic biclusters with the found biclusters, from which statistical evaluations such as true and false positive rates (TPR, FPR) and relative operating characteristics (ROC) curves are produced. **B.** Heatmaps of the gene-gene correlation matrices for all the synthetic data, the known synthetic biclusters (S1-8) and the biclusters found with FABIA (F1-8) and MCbiclust (M1-6). Numbers of gene and samples are shown in parenthesis (gene, sample) to compare the sizes of real biclusters with the ones found with either method.

### Analysis on E. coli Many Microbe Microarray database (M^3D^)

MCbiclust was applied to a extensive E. coli K-12 microarray data set from the Many Microbe Microarray database (M^3D^) (45). This dataset includes 907 samples and 7459 probes measured with Affymetrix microarrays and collated from a wide range of experimental setups from 39 different researchers, uniformly normalized using robust multi-array average (RMA). To find biologically relevant biclusters the MCbiclust pipeline was run 1000 times on random gene sets.

### Analysis on Cancer Cell Line Encylopedia (CCLE)

MCbiclust was applied to the CCLE dataset (46) composed of 969 samples with gene expression levels measured as mRNA using Affymetrix U133 plus 2.0 arrays and probe sets, again using uniform RMA normalization across all the samples. To study mitochondrial related biclusters, MCbiclust was run 1000 times on the 1098 MitoCarta (47) genes known to be related to mitochondria. MCbiclust was additionally run 1000 times on random gene sets containing 1000 genes to find biclusters affecting general pathways.

## RESULTS

### MCbiclust is uniquely designed to identify large biclusters

In order to validate MCbiclust and compare its performance with other biclustering methods, we have used a synthetic data set, modeling large biclusters, and a custom scoring system (See Materials and Methods and Fig. 2A). The dataset contained 8 known biclusters (on average a matrix of 130 samples and 500 genes), and 10 biclustering methods were tested (see Supplementary Table 1). Comparison of the known biclusters with the found biclusters was carried out as previously described ((36), see Fig. 2A). Based on these similarity analyses the quality of bicluster identification of each method was assessed. Table 1 shows the average true and false positive rates (TPR, FPR), precision, F1 score and a consensus score (36), taking into account the sum of the Jaccard index similarities of the predicted biclusters to their matched known biclusters, divided by the larger set. In addition, the consensus score includes a penalty for finding incorrect number of biclusters.

**Table 1.** Summary statistics for comparing the different biclustering methods. MCbiclust optimum refers to choosing the top samples and genes that maximise the Jaccard index to the known synthetic bicluster while threshold is the top samples and genes chosen from MCbiclust’s threshold method (see Supplementary Methods).

| Method | Biclusters Found | Consensus Score | Genes F1 | Samples F1 |
| --- | --- | --- | --- | --- |
| MCbiclust optimum | 6 | 0.4368 | 0.8145 | 0.6634 |
| MCbiclust threshold | 6 | 0.3462 | 0.8043 | 0.5864 |
| FABIA | 8 | 0.04106 | 0.1962 | 0.549 |
| FABIAS | 8 | 0.02475 | 0.2498 | 0.2878 |
| biMax | 8 | 0.002343 | 0.5697 | 0.01672 |
| CC | 8 | 0.0001895 | 0.02177 | 0.03344 |
| Plaid | 2 | 0.004164 | 0.1299 | 0.1747 |
| ISA | 504 | 0.001191 | 0.3256 | 0.5459 |
| FLOC | 8 | 0.0006008 | 0.06603 | 0.03746 |
| QUBIC | 9 | 0.0003819 | 0.008219 | 0.2113 |
| CPB | 24 | 0.0001685 | 0.02989 | 0.06277 |
| ISA best | 8 | 0.07504 | 0.3256 | 0.5459 |

MCbiclust has identified 6 out of 8 biclusters, and massively outscored the existing methods in precisely identifying large, so far hidden, large biclusters within the massive dataset. This includes outperforming FABIA, whose data model was used to design the synthetic data. ISA, which is designed to be used on large datasets, found over 500 biclusters, 8 of which were reasonable matches for the synthetic bicluster, thus it had a very large false positive rate, detecting small random biclusters. Even when considering only the correct 8 biclusters, ISA still had a lower performance than MCbiclust. For further evaluation of the different methods, we have plotted relative operating characteristics (ROC) curves for each synthetic bicluster. These results confirmed the higher sensitivity and specificity of MCbiclust compared to methods existing so far (see Supplementary Fig. S2).

Importantly, MCbiclust has an additional unique feature compared to existing methods. Apart from finding biclusters, it also ranks samples according to the strength of correlation between genes found in the bicluster. Principal component analysis can be thus further used to determine subclasses of samples in the ranking space. *PCA value versus ranking* plots revealed the distribution of the clustered samples in a characteristic fork pattern (Fig. 3A), probably indicating the polar distribution of samples along the average expression of the gene sets, responsible for the high correlation (see Fig. 3B, C, Supplementary Fig. S3 and Supplementary Methods).

**Figure 3.**
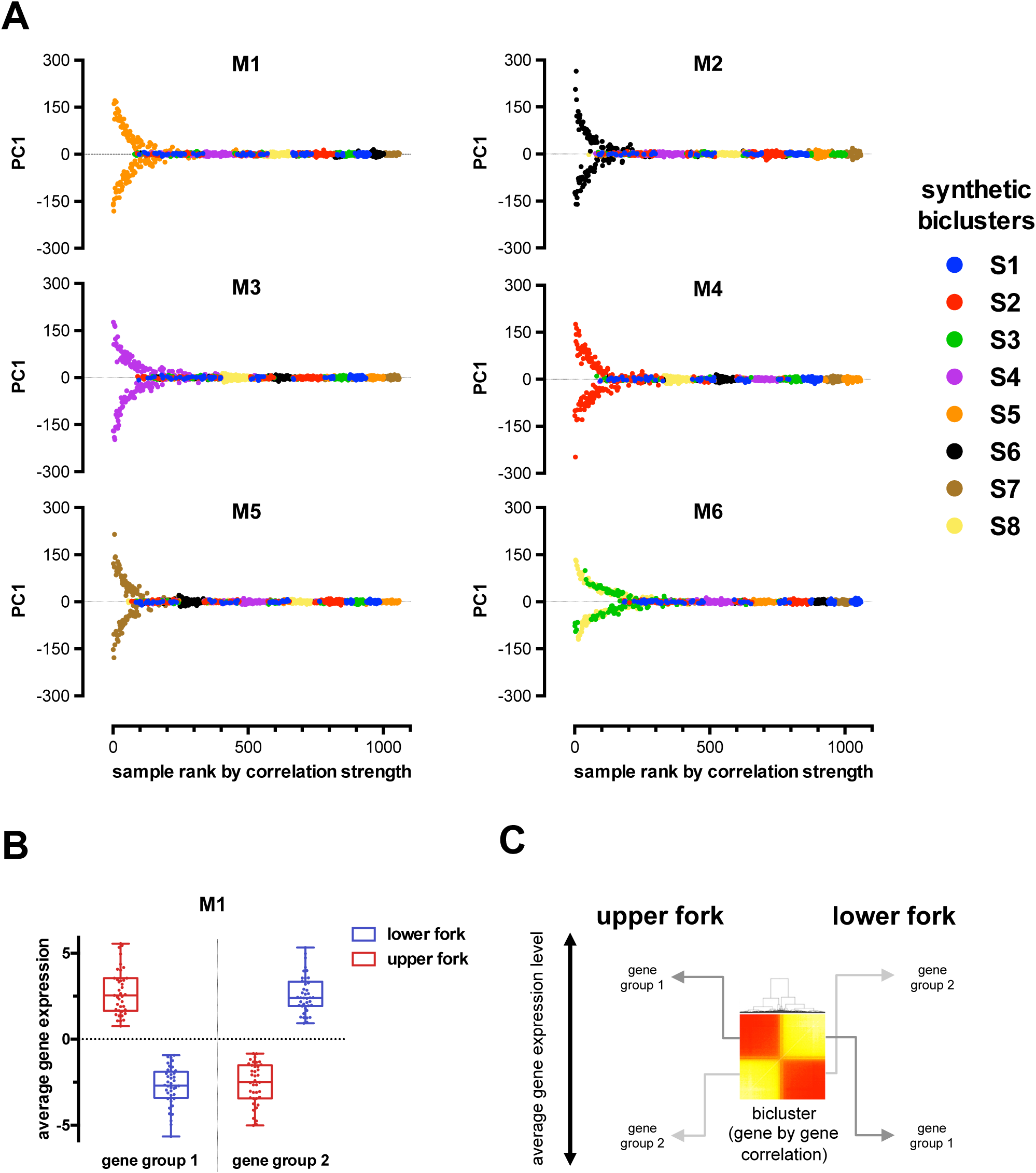
First principal component versus correlation based ranking plots of samples in biclusters identified by MCbiclust. Fork patterns of the six biclusters found with MCbiclust in the synthetic data. Y axes show the first principal component (PC1) value for each sample in each bicluster. Principal component analysis was run on the most highly correlating samples and captures the correlation pattern present in the samples. X axes show the ordering according to how well the samples preserve this correlation. Ranking is obtained as described in the ‘Extending the bicluster – samples’ section of Supplementary Methods. **B.** Mean centered average gene expression values of the two separate gene groups in the samples of the two forks of bicluster M1 determining the correlation. Expression levels in the two gene groups follow an antiparallel pattern. Relationship of average gene expression to PC1 values are shown in Supplementary Fig. S3. **C.** Schematics showing the gene-gene heatmaps of the M1 bicluster showing the division of the genes into two groups with different regulation in the upper and lower fork samples.

### MCbiclust discovers biologically relevant gene expression patterns in E. coli data sets

Next, we applied the algorithm to increasingly complex gene expression datasets from heterogeneous sample collections. First, we used an extensive E. coli K-12 microarray data set from the Many Microbe Microarray database (M^3D^) (45). The probes of this dataset cover ORFs or transcripts of unknown function as well as non-coding intergenic regions such as operon elements, 5’-UTRs, 3’-UTRs and small RNAs. The E. coli K-12 model is currently the best characterised prokaryotic model for studying gene regulatory networks on different scales, including large gene sets controlled by ! factors and smaller sets by transcriptional regulators. In addition, the dataset contains a large number of annotated experimental conditions, thus it was ideal for the initial characterization of MCbiclust’s ability to discover co-regulated gene sets in heterogeneous experimental conditions.

By running MCbiclust 1000 times, starting from random gene sets of 1000 genes, silhouette width analysis revealed 3 large distinct biclusters from the resulting correlation vectors (Fig. 4A, B). These groups were denoted E1, E2 and E3 and were obtained after 656, 229 and 115 runs, respectively, with the numbers indicating the runs required to reach dominance of the bicluster. These biclusters were all large; after thresholding with a sample p-value of 0.05 they contained 4822, 4700 and 6086 probes from 131, 130 and 96 samples, respectively.

**Figure 4.**
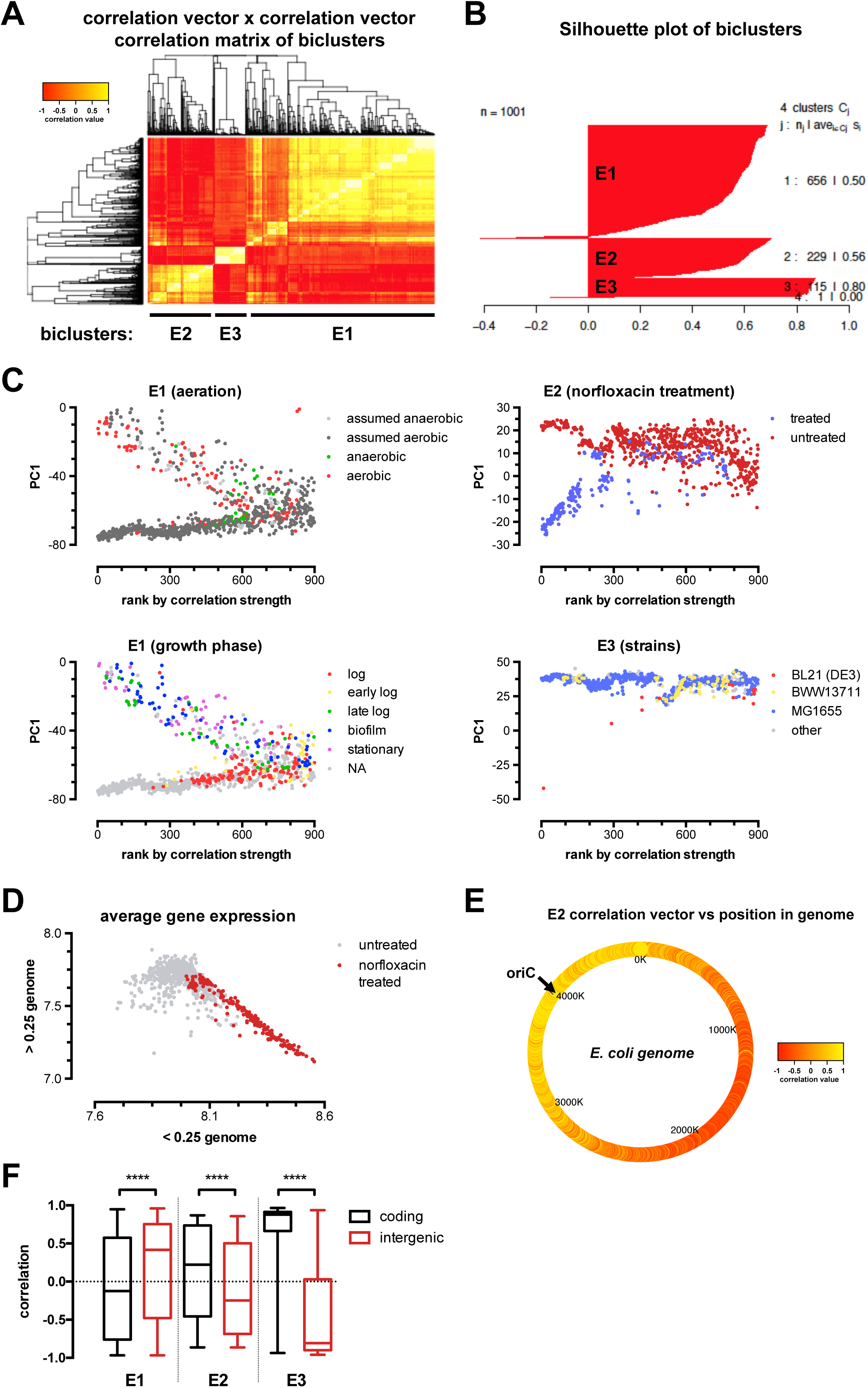
Biologically relevant biclusters discovered by MCBiclust in E. coli. **A.** MCbiclust was run 1000 times on the E. coli K-12 microarray data set from the Many Microbe Microarray database (M^3D^). Results are visualised in a heatmap of the correlation matrix from the correlation vectors. Hierarchical clustering reveals three large bicluster groups (E1-3). **B.** Correlation vectors are divided into three unique bicluster groups (E1-3) from the output of the silhouette analysis. The silhouette plot of the optimum number of clusters is shown as chosen by maximizing the average silhouette width of all the correlation vectors. **C.** PC1 versus sample ranking plots of the unique biclusters E1, E2 and E3. The plots have been overlaid with experimental conditions: aeration and growth phases for E1 (left panels), the gyrase inhibitor norfloxacin treatment for E2 (upper right panel) and the different strains used in the experiments for E3 (lower right panel). **D.** Plot of average gene expression values (median centered log_2_) close (<0.25 genome) versus far (>0.25 genome) to the origin of replication. The distribution of norfloxacin treated (red) and control (non treated, grey) samples are shown. **E.**Heatmap of correlation vector values for E2 in relation to genome position (oriC, origin of replication).**F.** Box plot of correlation vector values for all biclusters in coding (black) and intergenic (red) regions. The non-parametric Mann-Whitney test was used to calculate significance between pairs of each bicluster. **** p<0.0001

To understand the biological relevance of these biclusters, we first analysed the distribution of the samples in the found biclusters by PCA analysis and ranking according to the strength of correlation of gene expression (Fig 4C). As described above, the *PCA versus ranking* distribution plot typically gives a fork pattern, where the samples with highly correlated gene expressions are divided into high and low PC1 groups, where PC1 is mainly determined by the average expression level of the gene set defining the bicluster (see Supplementary Fig. S3). The plot allows the classification of the samples and helps to further determine correlations with sample types and experimental conditions. As shown in Fig. 4C, the samples identified in the E1 cluster were distributed along experimental conditions such as growth phase, aerobic/anaerobic status or treatment with antibiotics affecting growth. Cluster E2 clearly identified samples treated with a specific antibiotic, norfloxacin. In contrast, cluster E3 was determined by the highly deviant PC1 value associated with an outlier sample forming the upper fork of the distribution, while most of the samples remained in the lower half. Overall, the distribution analysis demonstrated the value of MCbiclust to identify biological (E1), pharmacological (E2) conditions, and outliers which otherwise would remain undetected (E3).

To identify more details of gene regulation in the biclusters we performed custom gene set enrichment analysis based on a Mann-Whitney test (see Supplementary Methods) to identify gene ontology (GO) terms related to E.coli, including Sigma factors and other E. coli transcription regulators from EcoCyc (48) and RegulonDB (49) databases. Additionally, terms for probes targeting either coding genes or intergenic regions were added. E1 and E3 had a large number of associated significant terms, 175 and 196, while E2 only had 25. Full tables of these terms are given in Supplementary Data. The custom analysis allowed the association of terms with positive and negative correlation vectors, informing on the average gene expression of pathways determining the distribution of samples in the upper or lower fork. The analysis revealed three important regulatory features.

First, the upper fork of E1 was driven by the correlated overexpression of genes with positive correlation vector values. Accordingly, those genes are predicted to drive an aerobic metabolic phenotype characteristic of slow growth in late log or stationary bacterial cultures or biofilms (see Fig. 4C). The terms cover wide range of metabolic pathways comprising biosynthetic routes of all major cellular components, lipids, proteins and ribonucleotide acids (see Supplementary Data), likely representing a specific global metabolic phenotype associated with the aerobic conditions in these experiments.

Second, the significant terms from E2 are relatively few and had relatively large p-values. Thus we looked at additional features of the genes determining the bicluster. Intriguingly, the average correlation vector values were distributed according to the position of genes in the E. coli genome (Fig. 4D). Indeed, Fig. 4E shows that this association can be explained by up-regulation of genes close to the origin of replication, which gradually decreased with the distance from the ORI. Examination of the conditions of the samples in this bicluster (see Fig. 4C) revealed that they have been grown in the presence of norfloxacin, a DNA gyrase inhibitor that prevents the division of the strands of E. coli DNA during replication, thus there would be two strands of DNA close to the ORI and a single strand further away, hence the gene dosage would be double around the ORI compared to genes further away resulting in this large-scale transcriptional difference in gene expression. Interestingly, a similar effect has been recently shown to exist in Streptococcus pneumonia by (50) but to our knowledge this is the first instance that reveal the effect in E. coli.

Finally, when we examined the terms which drive correlations in all three biclusters, the most significant associations were found with probes targeting either gene encoding or intergenic regions, which showed strong anti-correlation (Fig. 4F, Supplementary Data). Since average gene expression levels primarily determine PC1, our results show that expression of RNAs from intergenomic regions tend to exert inhibitory effects. This result is indicative of small non-coding regulatory RNAs that are intergenic inhibiting coding genes involved in biosynthetic processes and cell proliferation.

Altogether, MCbiclust therefore revealed three large-scale biologically relevant biclusters in the examined E.coli dataset: (i) one with terms linked to global metabolic changes during cellular growth in aerobic conditions, (ii) one showing how DNA gyrase targeting drug treatment stalls large-scale DNA replication and affects global gene expression and (iii) one that discovers a hidden sample preparation anomaly that seriously affects global gene expression in a single Affymetrix chip and possible other chips less severely (suggesting these chips should be removed before further analysis of this data collection). The results clearly indicate the value of MCbiclust to expose global trends in co-regulation of bacterial gene expression and other effects that cause changes in large-scale correlated gene expression within subsets of the biological samples.

### MCbiclust reveals cancer subtypes in the Cancer Cell Line Encylopedia data set

Next, in order to validate MCbiclust on highly complex and heterogeneous eukaryotic gene expression data, we have used a recently created cancer microarray dataset comprising ~1K cancer cell lines from diverse tissues of origin (Cancer Cell Line Encyclopedia, CCLE, (46)). Gene expression level heterogeneity between samples in this set arises from two main sources: (i) de-regulated gene expression triggered by the oncogenic genetic lesions and (ii) expression patterns distinctive of the tissue of origin of specific tumours. Here, due to the larger genome and sample numbers as compared to the E. coli dataset, we assumed that selection of the initial gene set might have substantial impact on the biclusters found and thus we have followed two different strategies. First, as described above we have run MCbiclust 1000 times utilising random gene sets, in order to discover potential large scale regulations affecting a subset of samples. In addition, however, we also sought to characterize specifically the regulation of multi-gene controlled global processes such as cellular metabolism and organelle biogenesis. Cancer evolution is known to involve radical rearrangements of cellular metabolism, in recent years deregulation of cellular energetics has even been recognised as an important hallmark of cancer (51). The aerobic glycolytic phenotype of many cancers for producing ATP has long been recognised, but it is less well understood how changes in mitochondrial biogenesis (here defined as co-regulation of the transcription of nuclear encoded mitochondrial genes, NEMGs) and hence energetic function affects cancer growth and survival. Thus our aim here was to investigate mitochondrial involvement in cancer using MCbiclust. Therefore, in the second instance MCbiclust was run on the CCLE dataset another 1000 times using a gene set composed of 1098 MitoCarta (47) genes, classified as NEMGs.

Silhouette analysis identified two distinct biclusters (R1 and R2) using random gene sets and one distinct bicluster (denoted M1) when using the MitoCarta gene set (see Fig. 5A and Supplementary Fig. S4). These biclusters can be directly compared by plotting the average correlation vectors of each measured gene in the genome between individual biclusters, as shown in Fig. 5B. Overall, we have found that the M1 and R2 biclusters are highly similar, with both having mitochondrial genes with high correlation values, thus both random and function-specific initial gene selection led to the identification of essentially the same bicluster.

**Figure 5.**
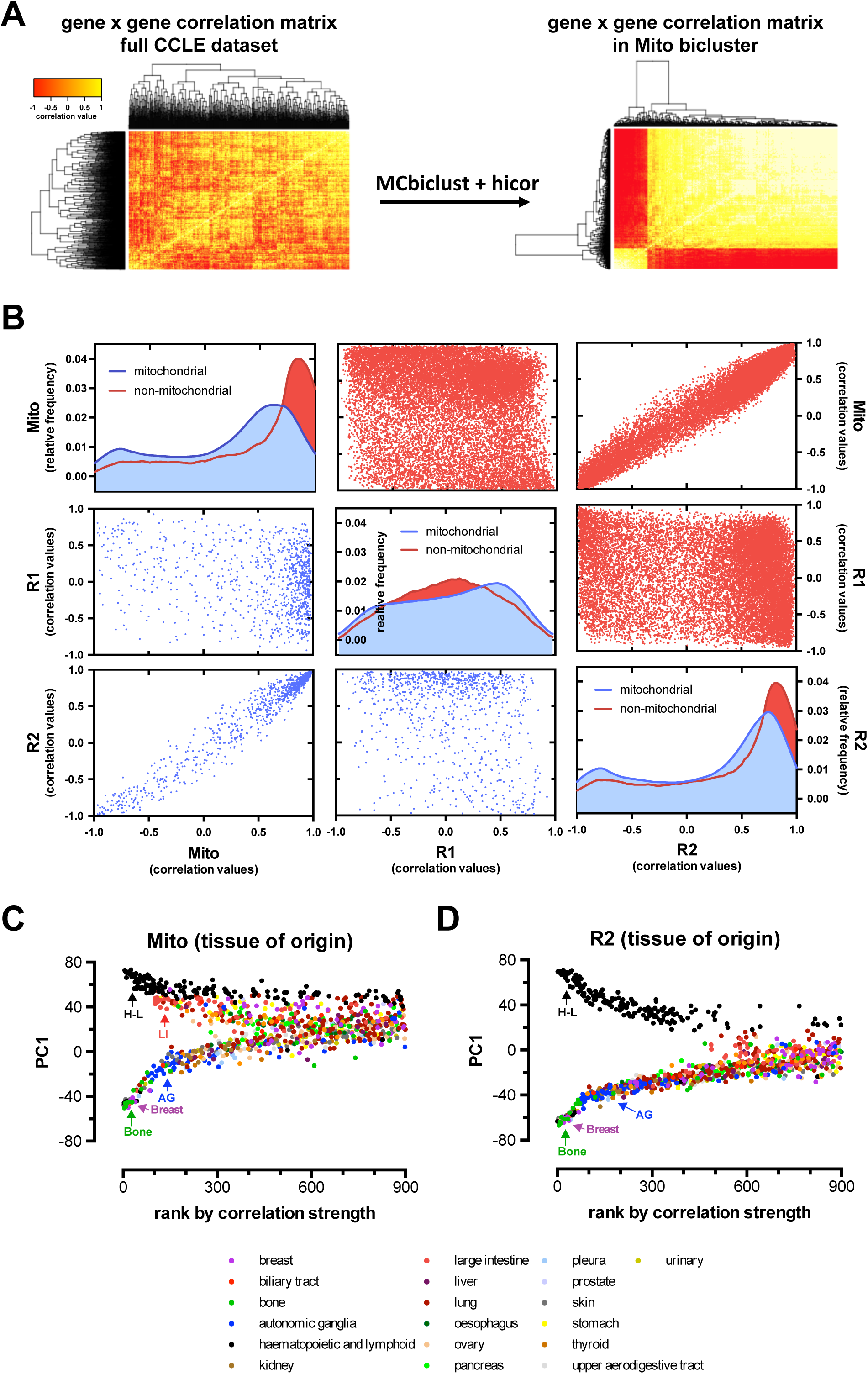
Biologically relevant biclusters in the Cancer Cell Line Encyclopedia (CCLE) microarray dataset. **A.** Heatmaps of the MitoCarta gene – gene correlation matrices across all the samples (left panel) and in the Mito bicluster of samples and genes established by the MCbiclust and Hicor algorithms (right panel), illustrating the biclustering process (see also Fig. 1 and Materials and Methods). Heatmaps and Silhouette plots of the distinct R1 and R2 biclusters identified using random initial gene sets are shown in Supplementary Fig. S4. **B.** A matrix of plots comparing the correlation vectors in all three distinct biclusters (Mito, R1 and R2). The diagonal plots show density histograms of the correlation values in the mitochondrial (blue) and non-mitochondrial gene sets (red) to the respective biclusters (Mito, upper left; R1 central; R2, lower right). Off-diagonal scatter plots show the relationships between the correlations of genes to the respective biclusters (Mito, R1 and R2, labelled left versus bottom) for mitochondrial (lower left triangle, blue) or non-mitochondrial genes (upper right triangle, red). **C.** PC1 versus sample ranking plots of the Mito and R2 biclusters, which are highly correlated (see scatter plots in panel **B**). The tissue of origin of the different sample cell lines is overlaid on the distribution plot. Clustered samples with the same tissue of origin are marked in the upper (Mito: H-L: hematopoietic and lymphoid, LI: large intestine; R2: H-L: hematopoietic and lymphoid) and lower (both Mito and R2: AG: autonomic ganglia, Breast, Bone) forks.

Next we performed the same custom gene set enrichment analysis (see Materials and Methods) done on each of the average correlation vectors as with the E. coli data. As shown in Supplementary data, the M1 and R2 biclusters define a functional group of genes highly related to the mitochondrial respiratory chain, but also ribosomes, ribosome biogenesis. This most likely represents activation of a novel gene regulatory pathway in a subset of samples (Fig. 5C, D), coupling increased mitochondrial biogenesis to cell growth. On the other hand, the R1 bicluster is highly enriched in immune system components and their regulated genes, and particularly overexpressed in a subset of carcinomas of different tissue origin (see Fig. 6A, B).

**Figure 6.**
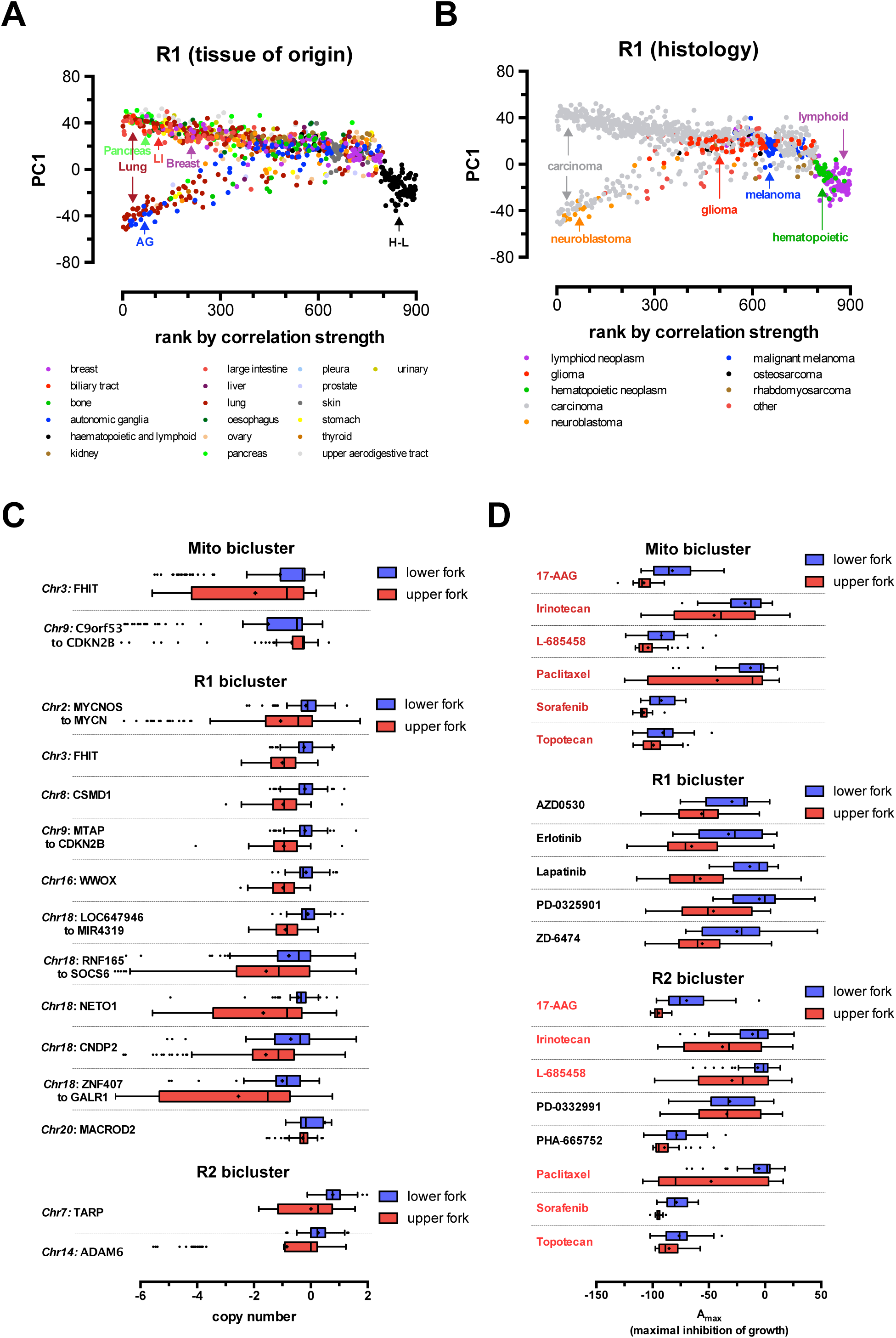
Pathological relevance of biclusters in the CCLE dataset. **A, B.** PC1 versus sample ranking plots of the R1 bicluster. The tissue of origin (**A**) and tumour histology (**B**) of the different sample cell lines is overlaid on the distribution plots. Clustered samples with the same tissue of origin or histology are marked across the distribution plots (LI: large intestine, AG: autonomic ganglia, H-L: hematopoietic and lymphoid origins). **C.** Association of copy-number differences across the whole genome with the distribution of samples in the upper and lower forks in all biclusters. Chromosome numbers and genes (labelled at left) with differences significant with a p-value < 0.05 are shown. **D.** Association of differences in pharmacological sensitivity to anticancer drugs with the distribution of samples in the upper and lower forks in all biclusters. To represent pharmacological sensitivity the A_max_ value was used from the CCLE dataset, signifying maximum inhibition of growth for each drug treatment. Drugs (out of 24 tested, see main text and ref) with significant differences between the lower and upper forks of each bicluster are shown. All differences are significant with a p-value <0.05. Significance in **C** and **D** was calculated using a permutation method randomly reassigning samples to the upper and lower fork and recalculating the average difference in copy-number or A_max_ values between the forks, and using this to form the distribution from which the p-values were calculated.

Finally, we further analysed the data to understand the potential association of the clustered gene expression patterns with the actual tissue of origin, pathology, genotype and pharmacological phenotype of the individual cancer cell lines. First, we mapped the relationship of the gene expression patterns of different cancer cell lines compared to the various biclusters. We ranked all samples according to the strength of correlations found in each bicluster, and plotted the rankings against the PC1 value for each sample. As shown above, PC1 values are mostly determined by the average gene expression values of a subgroup of genes in the bicluster (see Supplementary Fig. S3). Each bicluster was thus represented by the typical fork like distribution pattern (see Fig. 5C, D and Fig. 6A, B). This allowed us to overlay the tissue of origin and pathological subtype information on the distribution patterns. Whilst the mitochondrial M1 and R2 biclusters mainly separated cancer cell lines of hematopoietic origin from the rest of the tissues, the R1 bicluster had no tissue specificity. However, this bicluster was enriched in immune system related pathways and was typical to a subset of carcinomas (see Fig. 6A, B). Next, we calculated enrichment of locuses with gene copy number alterations (Fig. 6C) and pharmacological sensitivity to 24 anticancer drugs utilized in the CCLE study (46) (Fig. 6D). Importantly, various copy number alterations were found to be specifically associated with each bicluster, probably indicating the genetic, oncogenic origin of the gene expression patterns. Strikingly, the distribution between the upper and lower fork of the pattern also determined significant differences between the sensitivity to the growth inhibiting effects of various anticancer drugs in each bicluster (Fig. 6D), indicating the potential therapeutical predicting value of MCbiclust based cancer sample classification.

## DISCUSSION

MCbiclust outperforms other biclustering methods in terms of identifying large biclusters. The approach presented in this paper offers a new paradigm in the analysis of gene expression levels. This approach is pattern-centric, with large numbers of significantly co-regulated genes being sought unsupervised in a minority of the samples, once found both genes and samples can be ranked by how strongly an individual gene is being co-regulated in the pattern or how strong is this co-regulation in the sample. It has been demonstrated that the patterns it finds are biologically relevant and meaningful and it has great potential use in the analysis of transcriptomic datasets and classifying samples in a novel, biologically relevant way, according to their large scale gene transcription pattern.

A simple example for improving transcriptome analysis stems from the finding of a DNA replication effect hidden in the gene expression data within the M^3D^ E. coli data set (Fig. 4D, E). By revealing this effect, MCbiclust now makes it possible to normalise for it, e.g. in order to remove bias, allowing analysis of other gene sets with low signal strength.

Similar improvement in analysis can result from the finding in the third E3 bicluster. It is unusual in that a single sample with extreme global differences in gene expression has driven the formation of this bicluster. This sample was from an original study involving 16 Affymetrix arrays with 2 replicates over 8 conditions (52). Examining the images of the raw Affymetrix CEL files reveals that this sample (MGD1_t0_A.CEL) has very weak intensities over most of the chip compared to its replicate (MGD1_t0_B.CEL) and other samples within this study. This has probably arisen due to some problem with sample preparation since other aspects of the chip (such as spike-in concentration gradients) are normal. RMA normalization of this chip has brought these low gene expression values in line with other chips, but the normalization in turn causes a number of genes (mostly intragenic) to have abnormally high values. The resulting large-scale transcriptional pattern is what MCbiclust has detected within E3, and although not biological in nature, it does show the methods impressive power to find a single chip that has either sample or normalization issues within a very large data collection; thus potentially of use for data cleansing large –omics data collections. Interestingly Fig. 4C shows a few other samples within this data collection that potentially have similar sample preparation issues but not as extreme as this sample.

An intriguing feature of MCbiclust is that by creating *PC1 versus ranking* plots, the distribution and classification of samples can be better understood. Thus MCbiclust first discriminates samples according to the strength of correlation of a specific gene set, thus recognizes classes of samples with high and low correlation, indicating that a specific gene expression pattern is being regulated or not in in a specific class. However, since this regulation can be either positive or negative (creating anti-correlation patterns, see (Fig. 3), samples with higher expression of a subset of genes from the bicluster are clearly separated from samples having the gene set suppressed. This next level of classification, e.g. in the Mito bicluster, most probably reflects mitochondrial biogenesis (high in the upper fork samples), which is either activated or suppressed according to the metabolic needs of tumours (53). Such classifications have high chance of applicability both in discovery or clinical science based on gene expression data. For instance, since the correlation vector of the bicluster is known, expression of each gene of the genome, even outside the bicluster, can be correlated with it. Thus a correlation value can be associated which any gene, allowing the analysis of other cellular processes either acting upstream (e.g. master gene regulators of large gene sets or genetic changes), or downstream of the action of the bicluster. Of clinical relevance, correlation with clinical pathological phenotypes or as shown in Fig. 6C, D, differences in pharmacological sensitivity can be determined, thus probably allowing prediction of the phenotype of tumours. Interestingly, similar biclusters such as Mito and R1 in the CCLE dataset, predict slightly different tissue distribution (compare Fig.5C and D), indicating that the cellular phenotype is somewhat sensitive to small changes in the correlation vectors and the genes involved. Similarly, while the two biclusters predicted differential sensitivity between upper and lower fork samples to a common set of drugs (Fig. 6D), but bicluster specific drugs have also been found.

Another feature, and possible weakness in the current method is that a few biclusters can dominate the results which might exclude other biclusters to be found. Probably this is responsible for MCbiclust missing 2 synthetic biclusters (Fig. 2). By enriching the algorithm, we need to build an adapted version of MCbiclust that is more capable of identifying these weak signaled biclusters. In addition, apart from further developing the mathematical system, it will be of value to seek applications across all areas of gene expression research, from gene network regulation to biomarker discovery.

## ACKNOWLEDGEMENT

We are grateful for the helpful comments and discussions with members of the laboratory of GS participating in this project, in particular M. Menegollo, V. Pignataro and the University College London undergraduate students C. Quin, E. Tong, C. Esculier, S. Agarwal and Z. Ren.

